# The timescale and magnitude of 1/f aperiodic activity decrease with cortical depth in humans, macaques, and mice

**DOI:** 10.1101/2021.07.28.454235

**Authors:** Mila Halgren, Raphi Kang, Bradley Voytek, Istvan Ulbert, Daniel Fabo, Lorand Eross, Lucia Wittner, Joseph Madsen, Werner K Doyle, Orrin Devinsky, Eric Halgren, Mark T. Harnett, Sydney S. Cash

## Abstract

Cortical dynamics obey a 1/f power law, exhibiting an exponential decay of spectral power with increasing frequency. The slope and offset of this 1/f decay reflect the timescale and magnitude of aperiodic neural activity, respectively. These properties are tightly linked to cellular and circuit mechanisms (e.g. excitation:inhibition balance and firing rates) as well as cognitive processes (e.g. perception, memory, and state). However, the physiology underlying the 1/f power law in cortical dynamics is not well understood. Here, we compared laminar recordings from human, macaque and mouse cortex to evaluate how 1/f aperiodic dynamics vary across cortical layers and species. We report that 1/f slope is steepest in superficial layers and flattest in deep layers in each species. Additionally, the magnitude of this 1/f decay is greatest in superficial cortex and decreases with depth. We could account for both of these findings with a simple model in which superficial cortical transmembrane currents had longer time constants and greater densities than those in deeper layers. Together, our results provide novel insight into the organization of cortical dynamics, suggesting that the amplitude and time constant of local currents control circuit processing as a function of laminar depth. This may represent a general mechanism to facilitate appropriate integration of fast sensory inputs (infragranular) with slow feedback-type inputs (supragranular) across cortical areas and species.

## Introduction

Local field potentials (LFPs) in cortex exhibit pronounced oscillations of various frequencies(Buzsáki and Draguhn, 2004). Across the power-spectral-density (PSD) relationship, narrow-band oscillations such as alpha or gamma manifest as increases (‘bumps’) superimposed on a ramp of gradually decreasing power with increasing frequency (**Fig. 1c**). Across spatial scales, behavioral states and animals, this ramp is accurately modeled by an exponential function. This ramp of decreasing power, known as “1/f activity”, is aperiodic and conceptually distinct from rhythms like alpha or gamma, which are superimposed on it (**Fig. 1c**) (Gao, 2016).

**Figure 1:**
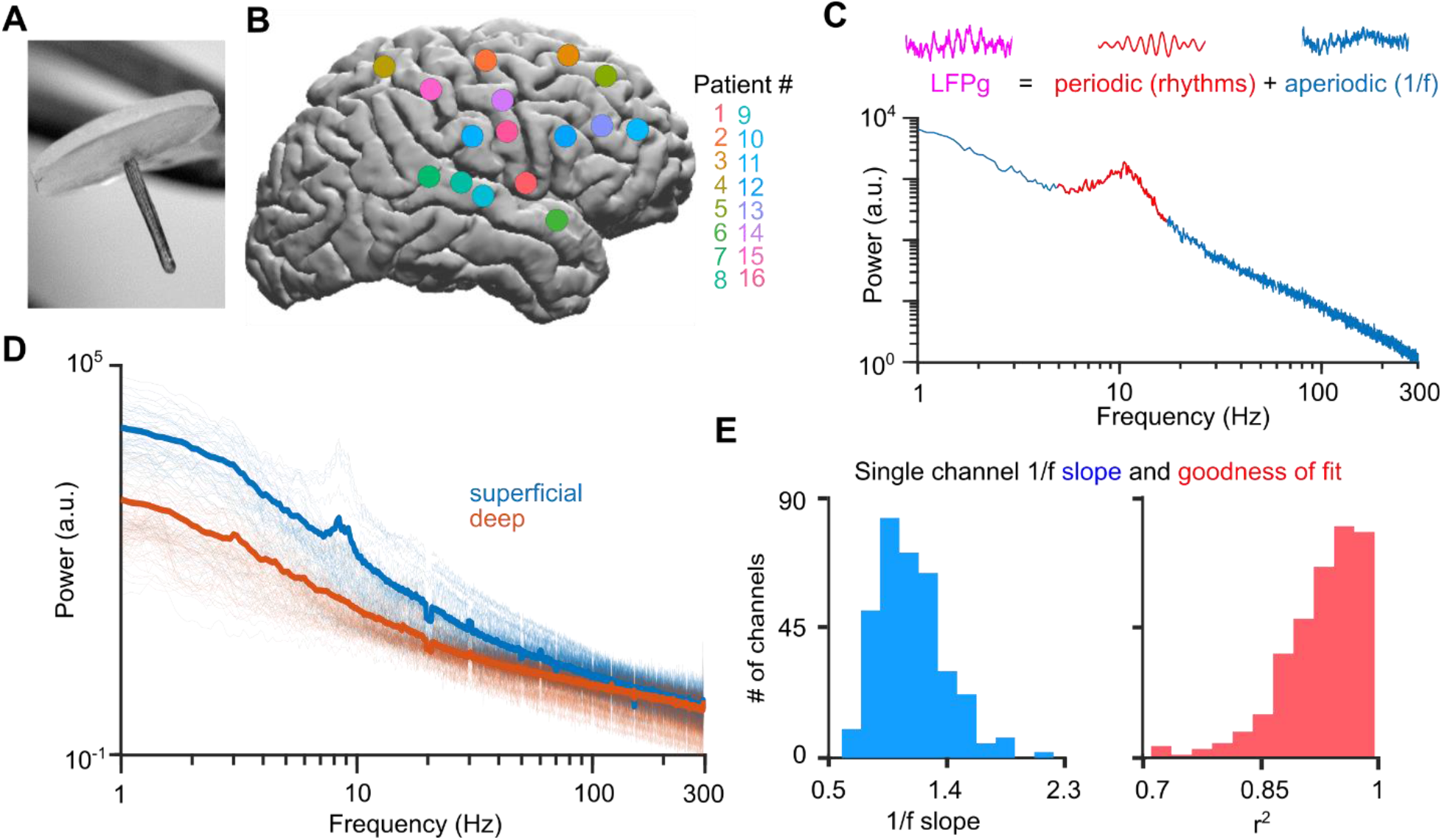
Laminar electrodes and aperiodic dynamics in humans. (**a)** Photomicrograph of a laminar array. Note the circular silastic sheet on top (adapted with permission from Ulbert et al., 2001). (**b)** Approximate locations of laminar implantations. We recorded from association cortex likely to be removed in prefrontal, temporal and parietal lobes. (**c)** Example wideband LFPg recording (purple trace) and power spectra (with 1/f fit) from Pt. 6, decomposed into periodic (red peak above 1/f line, red trace LFP bandpassed from 5-17 Hz) and aperiodic (blue power spectral density, blue time series bandstopped from 5-17 Hz) components. All LFPg traces are 1s. (**d)** All power spectra (light shade) and their means (dark) in superficial (blue, estimated layers I/II) and deep (orange, estimated layers V/VI) laminae. Superficial power spectra are both steeper and have greater offsets than deep ones. **(e)** Distribution of 1/f slopes and model fits to the 1/f.

1/f activity was previously disregarded as noise, but recent studies have shown that it’s slope and offset are correlated with cognition and behavior (Bódizs et al., 2021; Colombo et al., 2019; Freeman and Zhai, 2009; Gao et al., 2020; Lendner et al., 2020; Miller et al., 2009b; Ouyang et al., 2020; Podvalny et al., 2015; Waschke et al., 2021), age (Dave et al., 2018; Schaworonkow and Voytek, 2021; Voytek et al., 2015), pharmacological manipulation (Stock et al., 2019; Timmermann et al., 2019), and disease (Robertson et al., 2019; Veerakumar et al., 2019). Despite tracking such a broad range of biological and cognitive phenomena, the neural substrate(s) of 1/f activity remain unknown. Physiologically, 1/f may be a result of self-organized-criticality (Beggs and Plenz, 2003; V. Stewart and Plenz, 2006), population dynamics (Chaudhuri et al., 2018; Podvalny et al., 2015) or E-I balance (Gao et al., 2017).

1/f aperiodic dynamics can be quantified by a *P* ∝ *b* * *f*^−*α*^ power law, where *P* is power, *f* is frequency, *b* is a constant, and α is a positive exponent between .5 and 4 (He, 2014). Arithmetically, log *P* ∝ log *b* + log *f*^−*α*^, is the same as log *P* ∝ −α * log *f* + log *b*. Therefore, log power is linearly related to log-frequency (**Fig. 1c**). In this formulation, α is the slope of (log) power as a function of (log) frequency, and log *b* is its offset (though the PSD’s slope is negative, we refer to the slope α as positive by convention). The offset of this 1/f decay, or the power at which it intercepts the y-axis at *f* = 0 *Hz*, is proportional to the broadband amplitude of the LFP. Physiologically, this offset may correspond to mean firing rates or cortical activation (Manning et al., 2009; Miller et al., 2009a). The slope of this 1/f decay indicates a characteristic timescale, or memory, of the underlying signal (Milotti, 2002; Podvalny et al., 2015) (for the LFP, transmembrane currents (Buzsáki et al., 2012)). A slope of zero (flat 1/f) is the spectral representation of Poisson noise, and indicates no history-dependence; conversely, a large-magnitude slope (steep 1/f) indicates strong history dependence, or a long characteristic timescale **(Supplementary Fig. 1)**. This timescale reflects the degree of integrative processing in different areas: sensory cortex has a short timescale, while association cortex exhibits a long one. Certain behavioral tasks also exhibit different characteristic timescales (Gao et al., 2020).

Though previous studies have demonstrated that cortical LFPs obey a 1/f power law (Freeman and Zhai, 2009; He et al., 2010; Miller et al., 2009a; Voytek et al., 2015), none have measured whether 1/f dynamics change across cortical layers. This is important for understanding the generation of 1/f dynamics, as different cortical layers have distinct functional and anatomical properties and could be differentially responsible for generating aperiodic activity (He et al., 2010). Characterizing 1/f activity across layers could also tell us which laminae drive 1/f dynamics in scalp EEG and MEG. Insofar as the slope of the 1/f indicates a characteristic timescale of local processing (**Supplementary Fig. 1**) (Milotti, 2002), it may also indicate whether different cortical layers are specialized for integration over long periods versus fast non-integrative processing. To address this, we used microelectrode recordings made from humans, macaques and mice to measure the slope and offset of 1/f activity across cortical layers. We find that 1/f activity decreases in slope and offset with increasing cortical depth. Intriguingly, this relationship held across three mammalian species and multiple cortical areas.

## Results

To measure human cortical PSDs from different layers simultaneously, we made laminar micoelectrode recordings in 16 patients with medically intractable epilepsy (Halgren et al., 2018; István Ulbert et al., 2001) (**Fig. 1a-b**). During the implantation of clinical macroelectrodes, laminar microelectrode arrays were inserted into cortex expected to be resected. Each probe had 24 channels with 150 μm spacing, allowing us to record local field potentials throughout the cortical depth. To attenuate volume conduction, we referenced each contact to its neighbor, yielding the local-field-potential-gradient (LFPg) at 2000 Hz (filtered 0.2 – 500 Hz) (Halgren et al., 2018; Kajikawa and Schoeder, 2012; István Ulbert et al., 2001) (**Fig. 2a-b**). The LFPg yields a more focal measure of neural activity than the monopolar LFP without being as sensitive to noise as current-source-density(Trongnetrpunya et al., 2016). We analyzed data from epochs of quiet wakefulness, sleep, or when the state was unknown. When recordings from multiple states were available in a patient, we pooled data across states before further analysis, but patients in whom only sleep (Pts. 4, 5, 10) or wakefulness (Pt. 7) was recorded showed qualitatively similar effects. We then used Welch’s method to measure the Fourier Transform averaged across these epochs (10 second windows, single Hanning taper), giving us the PSD at each cortical depth (Halgren et al., 2018) (**Fig. 2c**). Though laminar probes span the cortical depth, the exact cortical layer can only be estimated based on previous measurements of laminar width(Hutsler et al., 2005).

**Figure 2:**
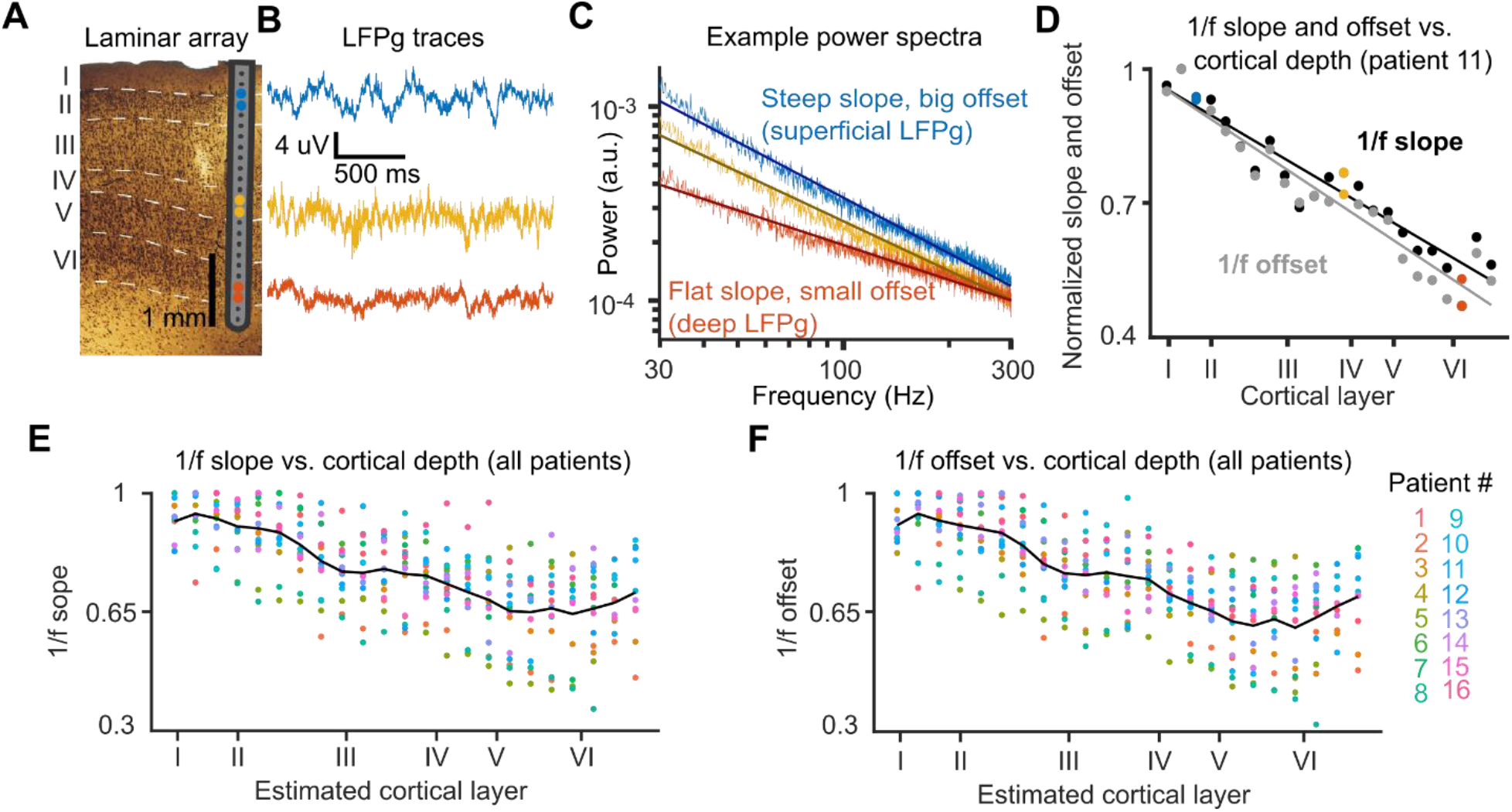
1/f slope fitting analysis in an example human patient. **(a)** Pt. 11’s laminar microelectrode array overlaid on a histological section of the tissue surrounding the probe; note that the array spans the gray matter. (**b)** Sample traces of the local-field-potential-gradient (i.e. the voltage difference between adjacent contacts) in superficial, middle and deep cortical layers. (**c)** 1/f power spectra (on a log-log scale) with fitted slopes from the spectral parameterization algorithm. Note that power spectra are well fit by a straight line, and that superficial channels (blue) have steeper slopes, whereas deep channels (orange) have flatter slopes. To emphasize this effect, we show these spectra from 30 – 290 Hz (though we fit on 1– 290 Hz). (**d)** The slope and offset of each channel’s power spectrum versus cortical depth in this participant. The slope flattens (i.e. becomes less negative) and offset decreases with increased depth – the colored dots come from the same sites as the colored traces and power spectra in (**b)** and (**c). (e)** 1/f slope and **(f**) offsets across all patients and contacts, normalized within patients.

Averaged PSDs from superficial layers (150 - 750 μm, layers I/II) exhibited both steeper slope and greater offset than deeper layers (2400-3450 μm, layers V/VI) (**Fig. 1d**). To quantify this, we fit slopes to the PSD for each channel using the spectral parameterization package (Donoghue et al., 2020). Briefly, the algorithm fits the aperiodic 1/f after removing Gaussian-fitted narrowband-oscillations which manifest as peaks on top of the 1/f. We fit the slope of each PSD from 1-290 Hz to capture spectral slope over a broad range. The spectral parameterization algorithm was effective at fitting our slopes, with an r^2^ across all channels of .93±.003 (**Fig. 1e**). Observed slopes were in the range of 0.60-2.21 (**Fig. 1e**), similar to previously reported values in human intracranial recordings (He et al., 2010; Lendner et al., 2020; Miller et al., 2009a; Voytek et al., 2015). However, comparing absolute slope and offset values across different recordings is problematic: differences in hardware and ambient noise can affect these measurements. Therefore, the most meaningful comparisons are of 1/f slope and offset within a recording (i.e. within patients) across channels. To implement this, we normalized all 1/f slopes and offsets within patients before visualization, such that the steepest (most negative) slope and largest offset had a value of 1 (**Fig. 2d**).

We first examined the relationship between 1/f slope and cortical depth within single patients. All 16 patients had a significant correlation (p < .05, uncorrected Pearson’s R) between cortical depth and 1/f slope; in all of these 16 recordings, 1/f slopes became flatter (less negative) in deeper layers, significantly more than expected by chance (p < .001, binomial test) (**Supplementary Fig. 2a**). The strength of this association varied, with r values ranging from -.95 to -.56 across patients, but on average (across patients) cortical depth explained nearly two-thirds of the variance of 1/f slope (r = -.78±.03, r^2^ = .62±.05, mean±standard error of the mean (SEM) across patients). This can also be seen clearly by pooling data across patients (**Fig. 2e)**. 1/f offset was also clearly related to cortical depth (**Fig. 2f**). Offset significantly decreased with cortical depth in all 16 patients (r = -.79±.03 across patients), similar to slope, indicating that broadband aperiodic activity is smaller in deep cortical layers (**Supplementary Fig. 3a**).

Were our results specific to humans or did they represent a general principle for mammalian cortical dynamics? To test this, we analyzed laminar recordings of spontaneous cortical activity in mice (Steinmetz et al., 2019) and macaques using Neuropixel probes. Mouse recordings were made from seven cortical areas, including visual, somatosensory, motor and retrosplenial cortex (**Fig. 3a, Table 1**); all macaque recordings were made from dorsal premotor cortex (PMd) (**Fig. 3b**). We selected experiments in which the probe was implanted approximately normal to the brain’s surface, and only analyzed contacts labeled as within the cortex. In order to make these recordings as comparable as possible to our human data, we spatially interpolated across channels to yield 23 bipolar LFPg series evenly spanning the cortical depth. Just as in humans, we used the spectral parametrization algorithm to quantify aperiodic slope and offset from 1-290 Hz with a strong goodness-of-fit (mouse: r^2^ = .92, macaque: r^2^ = .97 across single channels) and a plausible range of slopes (mouse: 0.81-2.55, macaque: 0.2-1.47). We found that 1/f slope (mouse: r = -.36 ± .11, macaque: r = -.88 ± .04 across experiments) and offset (mouse: r = -.47 ± .12, macaque: r = -.80 ± .08 across experiments) both decreased with cortical depth (**Fig. 3c-f, Supplementary Fig. 2b, 3b**, n=9 mice, 1 macaque).

**Table 1.**

| **Expt #** | **Cortical Area** | **Mouse** | **Session Label(s)** |
| --- | --- | --- | --- |
| 1 | Primary motor area | Moniz | Moniz_2017-05-18 |
| 2 | Primary motor area | Radnitz | Radnitz_2017-01-12 |
| 3 | Secondary motor area | Muller | Muller_2017-01-07 |
| 4 | Secondary motor area | Radnitz | Radnitz_2017-01-08,<br>Radnitz_2017-01-09,<br>Radnitz_2017-01-10 |
| 5 | Retrosplenial area | Lederberg | Lederberg_2017-12-07 |
| 6 | Retrosplenial area | Radnitz | Radnitz_2017-01-11 |
| 7 | Primary somatosensory area | Moniz | Moniz_2017-05-18 |
| 8 | Primary somatosensory area | Radnitz | Radnitz_2017-01-12 |
| 9 | Anterior visual area | Hench | Hench_2017-06-15 |
| 10 | Anterior visual area | Moniz | Moniz_2017-05-16 |
| 11 | Anterior visual area | Richards | Richards_2017-10-29 |
| 12 | Anteromedial visual area | Moniz | Moniz_2017-05-15 |
| 13 | Anteromedial visual area | Radnitz | Radnitz_2017-01-10 |
| 14 | Primary visual area | Cori | Cori_2016-12-14,<br>Cori_2016-12-18 |
| 15 | Primary visual area | Forssmann | Forssmann_2017-11-01 |
| 16 | Primary visual area | Hench | Hench_2017-06-15,<br>Hench_2017-06-17 |
| 17 | Primary visual area | Lederberg | Lederberg_2017-12-05 |
| 18 | Primary visual area | Moniz | Moniz_2017-05-16 |
| 19 | Primary visual area | Theiler | Theiler_2017-10-11 |

**Figure 3:**
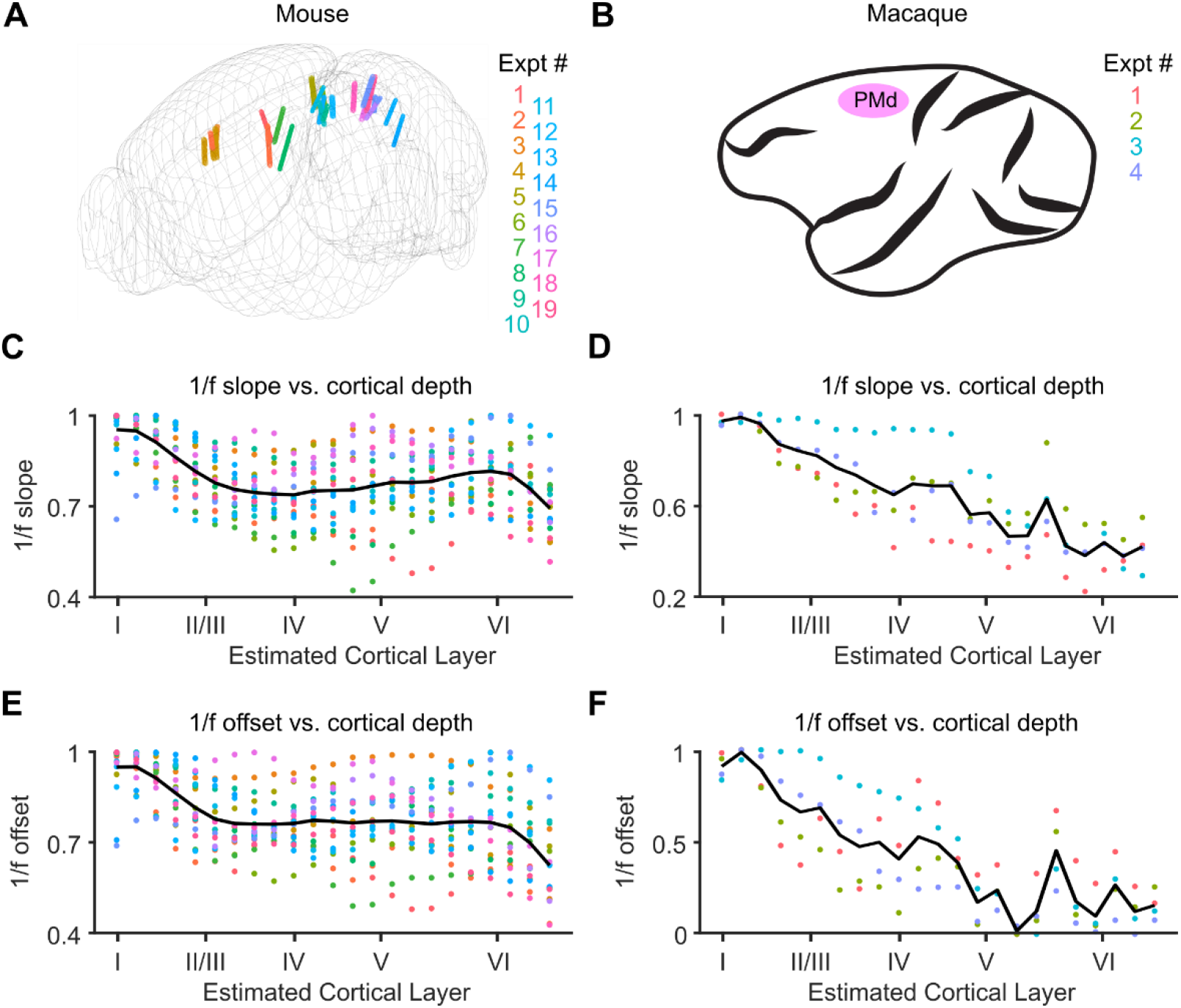
1/f fitting in macaque and mouse laminar recordings. (**a)** Approximate locations of Neuropixel probes. When multiple recordings were made from the same cortical area within the same mouse, the probes share the same color. (**b**) All macaque recordings were made from dorsal premotor cortex (PMd). **(c)** 1/f slope in mice and **(d)** macaques versus cortical depth, normalized within sessions. **(c)** 1/f slope in mice and **(d)** macaques versus cortical depth, normalized within sessions. **(e)** 1/f offset in mice and **(f)** macaques versus cortical depth, normalized within sessions (**c-f** as in **Fig. 2e-f)**.

Slope and offset were strongly correlated in each species (human: r^2^ = .94, p < 10^−10,^ mouse: r^2^ = .89, p < 10^−10^, macaque: r^2^ = .85, p < 10^−10^), even when depth was controlled for (human: r^2^ = .92, p < 10^−10^, mouse: r^2^ = .95, p < 10^−10^, macaque: r^2^ = .79, p < 10^−10^) (**Fig. 4b**). This strong relationship between slope and offset is mathematically equivalent to the power spectra of adjacent channels having a consistent x-intercept, and therefore “rotating” around a fixed frequency axis (**Fig. 4a**). If two power spectra intersect, a difference in their slopes will necessarily be accompanied by a proportional difference in their offsets (or vice-versa). Indeed, we found that throughout our datasets, the power spectra of adjacent channels consistently intersected at a similar median (human median: 106.77 Hz, mouse median: 115.41 Hz, macaque median: 28.68 Hz) (**Fig. 4c**).

**Figure 4:**
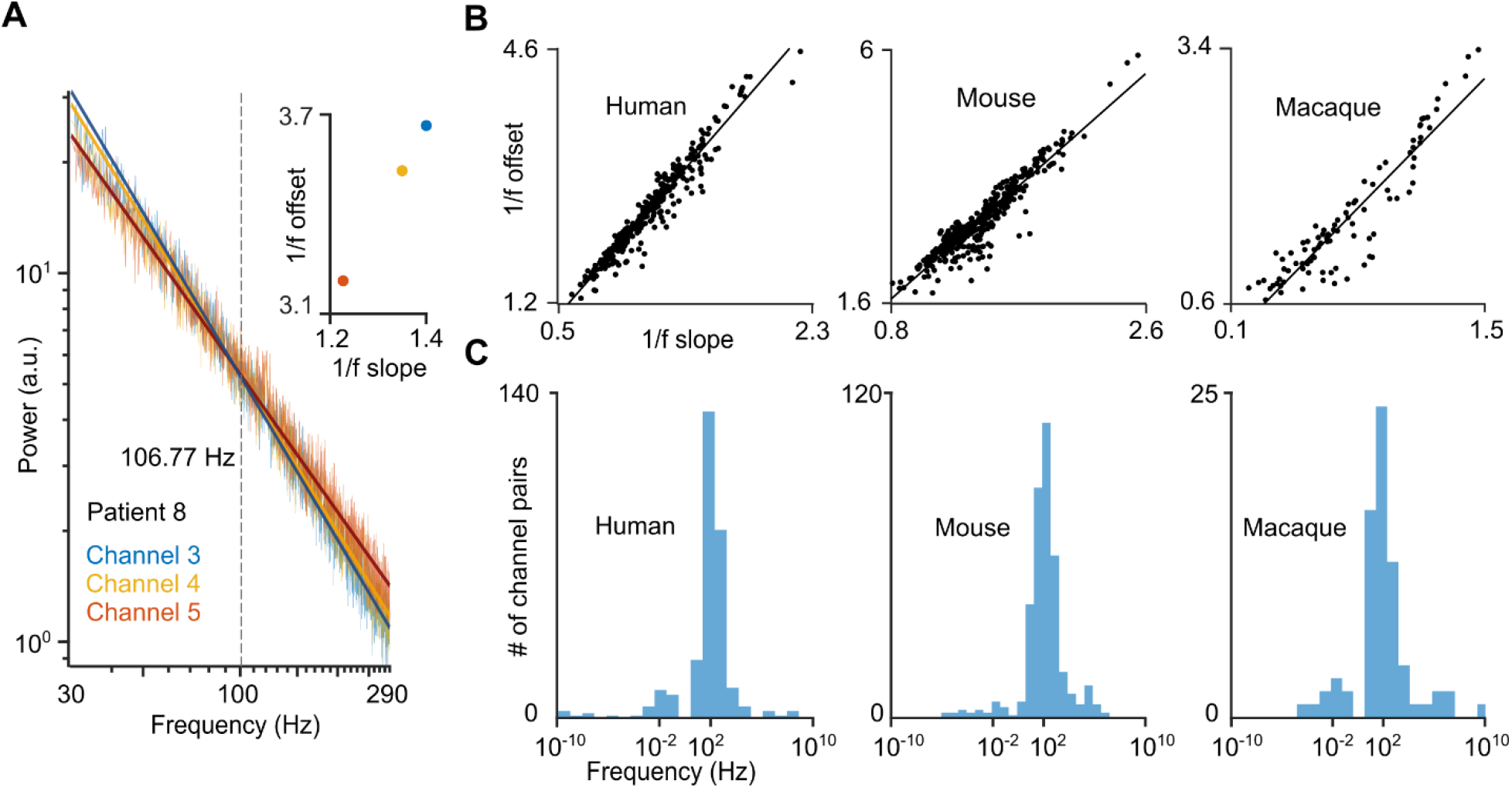
A common axis of rotation explains the correlation between 1/f slope and offset. **(a)** Three consecutive power spectra in an example patient; each of them intersect at approximately the same frequency of 106.77 Hz, the median intersection of all neighboring channels in humans. As seen in the inset, slope and offset are tightly correlated due to this common intersection. **(b)** 1/f slope versus offset for all single channels in each species. Note that 1/f slope and offset are highly correlated. (**c)** All intersection frequencies of neighboring power spectra across species; for visualization purposes, the 5^th^ percentile of highest and lowest intersection points in humans and mice, and the 10^th^ and 90^th^ in macaques, were excluded.

What might account for these robust differences in aperiodic dynamics across laminae? Several studies have shown that if the LFP is modelled by a Poisson spike train convolved with a postsynaptic conductance (PSG), and this PSG takes the form of a double-exponential, a 1/f power law emerges naturally (Freeman and Zhai, 2009; Gao et al., 2017; Miller et al., 2009a). This framework was used to show that a greater balance of inhibition over excitation (i.e. more GABA_A_ than AMPA PSGs) is associated with steeper 1/f slopes (Gao et al., 2017). Mathematically, this is due to the slower post-synaptic time constant of GABA_A_ compared to AMPA. Intuitively, a slower PSG will result in slower aperiodic dynamics, and therefore a steeper 1/f slope (as it has more low-frequency spectral power). A second discovery of these models is that the offset of 1/f power is directly related to the number of PSGs active; this has been previously used to argue that 1/f offset reflects mean firing rates (Miller et al., 2009a). Alternatively, differences in 1/f offset might reflect differences in postsynaptic receptor density (rather than differences in activation due to presynaptic firing). If each receptor is activated at an equal rate, an increase in receptor density should lead to an increase in postsynaptic currents and 1/f offset, or broadband LFP power. Accordingly, our results could be accounted for if: 1) aperiodic dynamics are affected by the time course of postsynaptic currents, 2) the ratio of postsynaptic channels activated with slow vs. fast time constants is largest in superficial layers, and 3) there is a greater receptor density in superficial than deep layers. We therefore hypothesized that a simple model which incorporates empirical receptor densities across layers in human cortex might reproduce the 1/f differences we observe with our laminar probes. Specifically, we utilized a database of receptor densities in 44 different cortical areas made from ex-vivo human tissue with autoradiography (Zilles and Palomero-Gallagher, 2017). Following previous work in modelling 1/f dynamics, we chose to model only the effects of ligand-gated AMPA and GABA_A_ receptors on the 1/f; these currents dominate excitatory and inhibitory postsynaptic currents in neocortex, respectively (Kandel et al., 1991). For each area, we generated two LFPs (and their PSDs) corresponding to supragranular and infragranular layers. Each LFP was generated by convolving Poisson spike trains (30 Hz) (**Fig. 5a**) with AMPA and GABA_A_ PSGs drawn from aggregating previous studies (**Fig. 5b**) (CNRG Lab @ UWaterloo, *Neurotransmitter Time Constants*). The number of spike trains convolved with AMPA and GABA_A_ PSGs was equal to the density of each receptor in fmol/mg within that layer.

**Figure 5:**
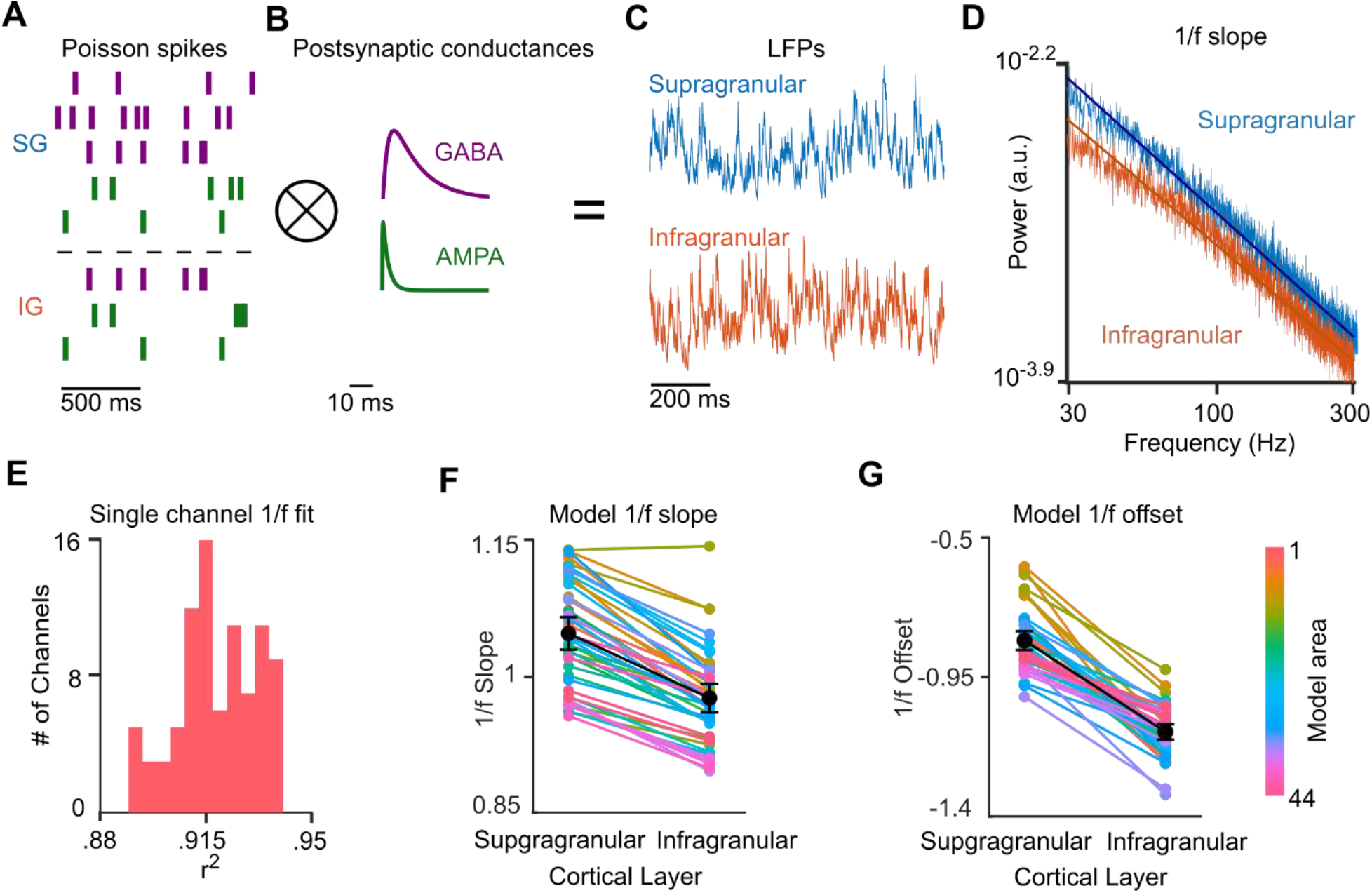
A simple model of 1/f dynamics across cortical layers based on AMPA and GABA_A_ receptor kinetics. (**a**) Poisson spikes corresponding to the fmol/mg density of GABA_A_ and AMPA in supragranular and infragranular layers are convolved with the (**b**) postsynaptic conductances (PSG) of these receptors. These yield LFPs (**c**) with log-linear power spectra (**d**), similar to our experimental recordings (see **Fig. 2c**). These spectra were then fit in log-log space with a straight line using the spectral parameterization algorithm (**e**). The slopes (**f**) and offsets (**g**) of these lines are higher in superficial than deep layers across areas.

Summing across these Poisson spikes yielded model LFPs for each layer and area (**Fig. 5c**). When we took the PSDs of these simulated LFPs (with the same window and taper we used on our actual laminar recordings), they were linear in log-log space (**Fig. 5d-e**) (goodness of linear log-log fit: r^2^ =.9980 for supragranular, r^2^ = .9968 for infragranular), and simulated 1/f slope was steeper in supragranular than infragranular layers for 43/44 model cortical areas (**Fig. 5f**) (p < 5.68*10^−14^ ; mean supragranular slope: -1.6243, mean infragranular slope: -1.4639, 7.29% difference in slope on average). 1/f offset was also greater in superficial than deep layers of all 44 model cortical areas (**Fig. 5g**) (p < 2.56 * 10^−12^; mean supragranular offset: -0.83, mean infragranular offset: -1.13, 25.92% difference in offset on average). These changes were comparable to our empirical changes in slope and offset between supragranular and infragranular cortex (22.48% rotation in slope, 25.41% change in offset). Similar to our empirical data, model 1/f slopes were highly correlated with offsets across simulated spectra (r^2^ = .49, p <= 3.49 * 10^−14^).

## Discussion

We find that aperiodic activity flattens and decreases with cortical depth across several cortical regions and three mammalian species. This suggests that the timescale and magnitude of aperiodic currents decrease with cortical depth as a general principle of neocortical activity.

Our data is suggestive of potential interspecies differences in aperiodic dynamics; for instance, the relationship between aperiodic dynamics and cortical depth was notable stronger in macaques and humans than in mice (compare **Fig. 2c-d** and **Fig. 3c-f)**. Unfortunately, differences in experimental conditions preclude direct comparisons between species. Distinctions in aperiodic dynamics across species could be due to different laminar probes (platinum-iridium vs. silicon), noise levels, or angles of insertion with respect to the cortical surface rather than physiological differences per se.

Using previously measured values of receptor density across human cortical layers in a simple model, we showed that the lower ratio of AMPA to GABA_A_ receptors in superficial layers, as well as the larger absolute number of receptors within superficial layers, could partially account for differences in 1/f slope and offset across cortical laminae. Insofar as the LFP reflects the activity of active postsynaptic channels, currents with slower timescales will lead to steeper slopes. Furthermore, the high density of receptors in superficial layers lead to greater 1/f offset (i.e. broadband power). Though several post-synaptic channels with long time constants such as NMDA are predominantly located in superficial layers (Eickhoff et al., 2007; Zilles and Palomero-Gallagher, 2017), we only modeled the effects of ligand-gated AMPA and GABA_A_ receptors on the LFPg, similar to other recent models of 1/f activity The strong correlation of 1/f slope and offset suggests a common physiological factor; our model indicates this could be due to a high density of postsynaptic channels with long time-constants in superficial laminae. However, it is important to note that other (non-exclusive) factors also likely contribute to laminar differences in aperiodic dynamics. One other such factor includes voltage gated (active) channels. Specifically, h-currents may contribute to the steep slopes / slow aperiodic dynamics observed in supragranular laminae. Firstly, h-currents can strongly shape the extracellular LFP (Ness et al., 2018). Secondly, the density of h-channels on the dendrites of layer 5 cortical pyramidal cells increases with distance on the apical trunk from the soma (Harnett et al., 2015; Kole et al., 2006), mirroring the smooth increase in 1/f slope from deep to superficial cortex. Additionally, human supragranular cells exhibit higher densities of h-channels when compared to rodents (Beaulieu-Laroche et al., 2018; Kalmbach et al., 2018); this may explain why the correlation between 1/f slope and depth was stronger in primates than mice. Alternatively, our effects could be due to biophysical differences across cortical layers. For instance, a gradual change in cortical impedance across layers could lead to different 1/f slopes and offsets, though cortical impedance is isotropic in macaque cortex (Logothetis et al., 2007). Similarly, gray matter may intrinsically filter extracellular currents in a frequency-dependent manner (though this is highly contentious (Bédard et al., 2006; Bédard and Destexhe, 2009)) in a way that changes between layers. These other factors may be incorporated in future, more biophysically complete models.

We employed a bipolar referencing scheme to emphasize local activity (**Fig. 2a**). Monopolar LFP recordings, made with a distant reference, may show an entirely different relationship between 1/f dynamics and cortical depth (Shirhatti et al., 2016), likely due to contamination by volume conduction (Kajikawa and Schoeder, 2012). Because infragranular LFPs are particularly susceptible to volume conduction from superficial cortex (Kajikawa and Schroeder, 2015), a monopolar reference might even find the opposite of our results (i.e. 1/f slope and offset would increase with cortical depth), despite the neural generators of these aperiodic dynamics residing in superficial layers.

Previous work has shown that currents in superficial layers underlie low-frequency oscillations, which are distinct from the aperiodic dynamics reflected in 1/f slope (Cash et al., 2009; Csercsa et al., 2010; Haegens et al., 2015; Halgren et al., 2015, 2019, 2018) (**Fig. 1c**). Crucially, the spectral parameterization algorithm is able to remove the oscillatory peaks before fitting the slope and offset of our power spectra. Furthermore, our results were robust even when slopes were fit from 30-290 Hz, far outside the range of delta, theta, alpha or beta oscillations (**Supplementary Fig. 5**). Therefore, both slow oscillatory activity and slow aperiodic dynamics are concentrated in superficial layers. It’s possible that channels with long time constants (such as HCN or NMDA receptors) are responsible for sustaining both low-frequency oscillations and slow aperiodic dynamics. Whether this implies a common physiological origin should be explored in future work.

What do differences in 1/f slope between cortical layers imply about the functional role of different laminae? 1/f represents the history-dependence, or “memory” of the LFPg (**Supplementary Fig. 1)** (Milotti, 2002). For cognition, the brain must simultaneously represent multiple timescales at different orders of magnitude(Kiebel et al., 2008). Indeed, it’s been previously shown that sensory cortex has the shortest processing timescales (presumably for encoding veridical representations of fast-changing stimuli), and that association cortex has the longest (Runyan et al., 2017). These timescales (as measured both by the LFPg and spiking) smoothly increase from lower to higher order cortex (Gao et al., 2020; Murray et al., 2014; Siegle et al., 2021; Spitmaan et al., 2020). Our work indicates that the gradation of neuronal timescales across areas is mirrored by differences in timescales across layers: currents in superficial layers have a longer timescale than currents in deep layers. This is anatomically expected, as superficial layers are the primary recipients of feedback and modulatory inputs (thought to have long timescales), whereas deeper layers receive more feedforward, driving inputs (thought to have a short timescales) (Markov et al., 2014). The long timescale of supragranular currents could allow them to have stronger integrative properties than infragranular currents, which may be specialized for faster bottom-up inputs.

## Materials and Methods

### Human Laminar Recordings

Laminar microelectrode arrays were inserted on the basis of two criteria: first, the tissue must be very likely to be resected (Istvan Ulbert et al., 2001). This could be because it was clearly within the seizure onset zone, or because it was healthy tissue overlying the seizure onset zone which would have to be removed during the resection. Secondly, the cortex in question must have had no chance of being eloquent. A silicone sheet attached to the array’s top was used to keep the probe perpendicular to the cortical depth, with surface tension between the sheet and the pia, as well as pressure from the overlying grid and dura, keeping the array in place. This sheet also ensured that the laminar array was perpendicular to the cortex and that the first contact was placed ∼150 microns below the cortical surface. Electrode impedances and phases were measured prior to implantation to ensure that recording properties were similar across contacts. Each laminar probe spanned the cortical depth with a length of 3.5 mm and diameter of .35 mm. Contacts had a 40-micron diameter spaced every 150 microns. Recordings were made during for an average of 17.32 minutes of task-free activity in each patient.

In patients 9 and 11, co-histology was performed on the tissue surrounding the laminar probe to confirm which layers each channel recorded from (**Fig. 2a**). In other patients, the correspondence of channels to individual laminae was approximated from previous measurements of laminar width. Channels 1, 4, 9, 14, 16 and 21 were the approximate centers of layers I-VI, respectively.

### Human clinical implantation details

All data were visually inspected for movement, pulsation and machine artifacts. The data was also screened for epileptic activity such as interictal discharges and pathological delta by a board certified electroencephalographer. Laminar arrays with significant amounts of artifactual or epileptiform activity, and/or insufficient technical quality, were rejected prior to further analysis. All epochs with artefactual or epileptiform activity from accepted arrays were also excluded from analyses.

The effects of Anti-Epileptic Drugs (AED) on the LFP is a potential concern, as these sometimes affect scalp EEG (Blume, 2006). While some patients in this study may have been taking AEDs, most recordings were performed after the patient’s medications had been tapered to encourage spontaneous seizure occurrences during the monitoring period. Critically, the results were highly consistent across all participants regardless of medication history, etiology, electrode location, or degree of epileptic activity. Expert screening and consistency across participants strongly suggest that our findings are generalizable to the non-epileptic population.

Because we did not systematically track behavioral state with EMG or sleep-scoring from scalp EEG, we were not able to investigate the effect of state on 1/f dynamics. However, in a subset of patients recordings were made only during definitive wakefulness (Pt. 7) or sleep (Pts. 4, 5, 10). The single-subject relationship between 1/f slope and offset with depth was highly similar in all cases (**Supplementary Fig. 2a, 3a**), suggesting that our findings generalize across states.

### Mouse laminar recordings

Mouse recordings were used from “Distributed coding of choice, action, and engagement across the mouse brain”(Steinmetz et al., 2019). Details on these experiments, as well as all of the data we analyzed, can be found at (https://figshare.com/articles/Dataset_from_Steinmetz_et_al_2019/9598406). Only implantations which were approximately normal to the cortical surface were selected for further analysis, to avoid confounding cortical depth with lateral or anterior/posterior position. This was done by visual inspection of electrode trajectories on the Allen Institute common-coordinate-framework(Wang et al., 2020). However, because these recordings were not made with this in mind, these recordings were likely not as normal as our human recordings (**Fig. 3a**). Mice were headfixed to perform a visual contrast detection task. We analyzed both periods of spontaneous activity when the mouse was neither being presented stimuli nor actively performing a task (spontaneous.intervals.npy). When multiple sessions from the same cortical area in the same mouse were available, we averaged spectra across all available sessions.

Only channels labeled as being in cortex were analyzed. We first spatially averaged Neuropixel LFPs between consecutive blocks of four adjacent channels (yielding 96 pseudochannels per probe) to reduce noise. Then, a spatial Gaussian filter with a standard deviation of 68 um was applied. To reject superficial contacts which were out of the brain, we only analyzed pseudochannels below or the pseudochannel immediately above the pseudochannel in which the most superficial unit was recorded. To make the spatial resolution of our recordings similar to humans, we then spatially interpolated evenly across the pseudochannels to get LFP timeseries spanning the cortical depth. Finally, we took the difference between these adjacent channels to derive the 23 channel mouse LFPg.

### Macaque Laminar Recordings

Rhesus Neuropixels recordings (4 sessions, 1 macaque) were performed in dorsal premotor cortex (PMd) using Neuropixels NHP probes supplied by IMEC and Janelia. We analyzed an average of 37.42 minutes of task-free, spontaneous activity per session. The Neuropixels probe was inserted along the axis of the recording chamber with the intent to be orthogonal to the surface of the cortex and spanning cortical lamina. Neuropixels probes were inserted using a custom adapter to mount the probe on a microelectrode drive (Narishige). A blunt 23-gauge stainless steel guide tube was mounted in a second tower and positioned in contact with the surface of the dura. The granulation tissue and dura was perforated using a tungsten electrode, then the Neuropixels probe was inserted through the same guide tube into the brain. Recordings were performed using the 384 channels closest to the probe tip, spanning 3.84 mm of length along the probe. The recording depth was selected so as to situate the most superficial observable spiking neurons aligned with the top recording channel. LFP data were acquired using SpikeGLX software and were digitized at 1kHz. Before fitting these LFPs, we applied the same procedure to our macaque recordings that we applied to our mouse recordings (i.e. spatially averaging adjacent channels, interpolating, and then taking the potential difference between these pseudochannels).

### Analysis

All analysis was performed in MATLAB (R2019b, Natick, MA) using custom and FieldTrip functions (Oostenveld et al., 2011). In each participant, the Fourier Transform was calculated in 10 second epochs on the zero-meaned data after a single Hanning taper was applied. FFTs were taken across all recordings made in each patient, even if recordings were made during multiple behavioral states, yielding a single 1/f slope at each cortical depth per patient. The frequency spectra of faulty channels (an average of 3 per probe) were linearly interpolated from the normalized frequency spectra of good channels above and below on the laminar probe. For instance, if channel 2 was defective, its power spectrum would be replaced by the average of channels 1 and 3’s power spectra. To ensure that interpolation wasn’t yielding spurious results, we examined the 1/f depth profiles of patients without any interpolated channels (2, 3, 7, 8, 9). These patients displayed the same pattern as those with interpolated channels.

Power spectra were visually examined for noise peaks, such as 60 Hz line noise, and power values at ±.5 Hz around these high-frequency noise artifacts were removed. After this, we normalized the power spectra within each patient by the median power value across all channels and frequencies from 1-290 Hz (within that patient). In both humans and mice, we fit on all frequencies (excluding artifactual noise peaks) from 1 – 290 Hz. In humans, we used a peak sensitivity rating of 7, and in mice we used a peak sensitivity rating of 4. Peak width limits of 3-14 Hz were used in both species.

As a first control analysis, we replicated our results using a simple linear fit with a first-degree polynomial (polyfit.m) after both power spectra and the frequency axis were log transformed. This linear fit (in log-log space) found the same relationship between slope/offset and cortical depth (**Supplementary Fig. 6**), demonstrating that our results are robust to different fitting algorithms.

Another concern is that a correlation between the spectral parameterization algorithm’s goodness of fit (r^2^) with cortical depth, offsets or slopes could yield spurious relationships between depth and offset or slope and offset. To ensure that this was not the case, we measured the Pearson correlation between goodness of fit and depth, slope and offset across all channels within our mouse and human datasets (separately). All of these correlations in humans were insignificant (r^2^ <= .006, p >= .14), and were very weakly correlated (albeit significantly) in mice (r^2^ <= .06, p >= 1.05 * 10^−7^). We there for re-measured the average correlation between slope/offset and depth for only channels in a very narrow range of goodness of fits (r^2^ = .99-1), reducing the correlation between goodness of fit and depth, slope or offset. Even with this restricted range of channels, we observed high correlations of slope and offset to cortical depth (r^2^ = .31, .48 for slope and offset respectively). This suggests that our results cannot be explained by systematic variations in our fitting accuracy.

To make sure our slope-fitting wasn’t contaminated by low-frequency oscillatory peaks (though these should be removed by the spectral parameterization algorithm), we replicated our results using a fitting band of 30 – 290 Hz. 30 Hz was chosen as a lower bound to exclude delta, theta, alpha and beta oscillations (**Supplementary Fig. 5**).

As a last control, we re-ran our fitting across a wide range of frequency bands. 1/f slope/offset and cortical depth were strongly correlated across almost all of these control bands (**Supplementary Fig. 4**). That these correlations remained even when the upper frequency bound was relatively low (<80 Hz) suggest that our results are not due to residual spiking contamination of the LFPg.

A point of potential conceptual confusion is our use of the aperiodic 1/f exponent, as opposed to the knee-frequency, to derive timescale. Though some previous papers have used the knee-frequency to measure the timescale (Gao et al., 2020), we rarely found spectral knees in our recordings. Furthermore, both the aperiodic exponent and spectral knee (usually not present in our data) determine LFPg timescale (Gao et al., 2020; Milotti, 2002) (**Supplementary Fig. 1**). This justifies our use of the aperiodic exponent instead of the knee-frequency for our analysis.

### Model

Our model is similar to earlier studies of 1/f dynamics (Freeman and Zhai, 2009; Gao et al., 2017; Miller et al., 2009a). We modelled LFPs as a convolution between Poisson spiking (30 Hz for 490 seconds) and post-synaptic conductance kernels for AMPA and GABA_A_ receptors. The number of Poisson spikes convolved with each receptor-type was equal to the (rounded) density in fmol/mg of that receptor. We then summed these convolutions within the supragranular/infragranular layers of each cortical area, yielding 88 artificial LFPS (Zilles and Palomero-Gallagher, 2017). The FFT was made using the same window and Hanning taper as our laminar recordings, and plotted in log-log space. Slopes were fitted in the same way as our empirical laminar recordings in humans (i.e. with the spectral parameterization algorithm from 1 – 290 Hz, peak sensitivity of 7 and peak width limits of 3-14 Hz).

### Code Accessibility

Data and analysis scripts are freely available (https://github.com/harnett/LaminarAperiodic)

## Acknowledgements

We thank the patients for participating in this research and Eric Trautmann for the generous contribution of macaque laminar recordings, as well as Thomas Donoghue, Norma J. Brown, Scott Cole, Enrique Toloza, Adam Niese, Dimitra Vardalaki and Ian Weaver for constructive feedback and technical support.

This work was supported by the Harvard MIT/Joint Research Grants Program in Basic Neuroscience and Tiny Blue Dot Foundation (M.T.H. & S.S.C.), as well as National Institutes of Health Grants R01-MH-099645, R01-EB-009282, R01-NS-062092, K24-NS-088568, the U.S. Office of Naval Research Grant N00014-13-1-0672 (E.H. & S.S.C), the MGH Executive Council on Research (S.S.C.), Hungarian National Brain Research Program grant KTIA_13_NAP-A-IV/1-4,6, EU FP7 600925 NeuroSeeker, and Hungarian Government grants KTIA-NAP 13-1-2013-0001, OTKA PD101754 (I.U., D.F., L.E., L.W.). M.T.H is a Klingenstein-Simons Fellow, a Vallee Foundation Scholar, and a McKnight Scholar.

---

**Supplementary Figure 1:**
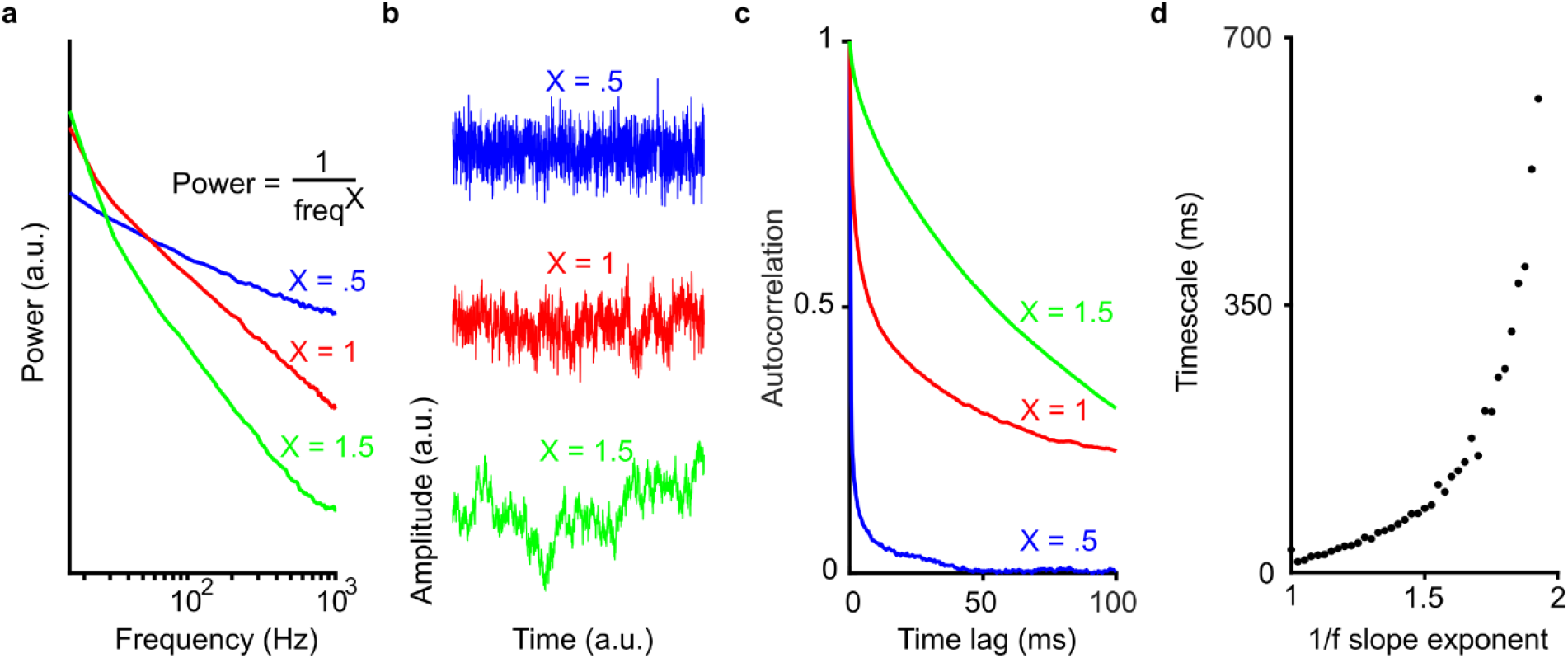
1/f slope reflects timescale of activity in the time-domain. **a)** Three simulated power spectra with different slope exponents (.5, 1, 1.5). (**b**) 1 s of simulated time series corresponding to the spectra in a. Note that the time series with the slowest fluctuations corresponds to the spectrum with the steepest exponent. (**c)** Autocorrelation functions for the three simulated timeseries in **b**. The highest autocorrelation function (i.e. most time dependence or memory) corresponds to the steepest power spectrum (X = 1.5). **d**) Timescale as a function of slope exponent for 38 simulated time series (slopes between 1 and 2). Timescale is defined as the lag at which the autocorrelation decays to 1/e.

**Supplementary Figure 2:**
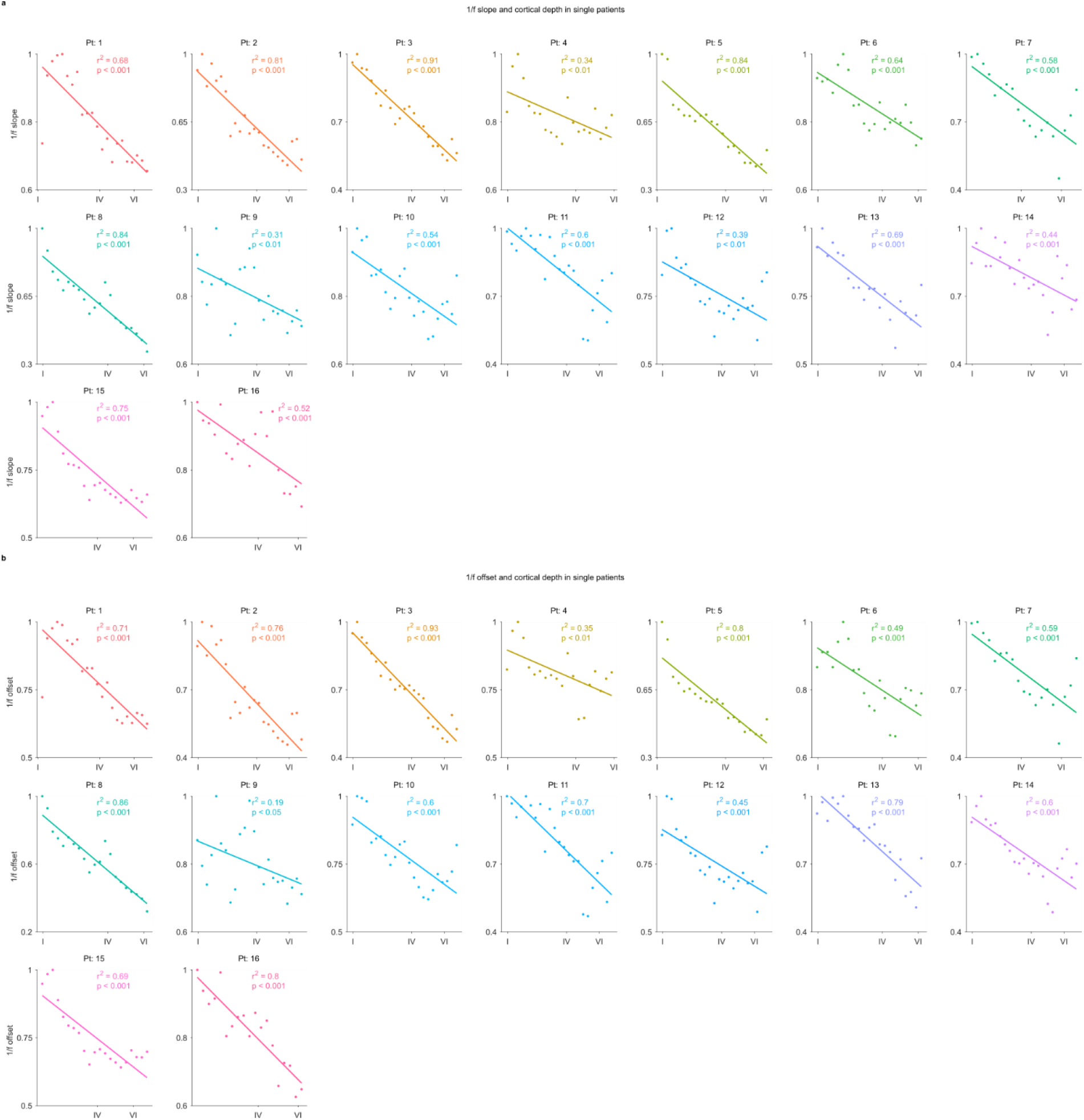
1/f slope and offset decrease with cortical depth in single patients. 1/f slope (**a)** and offset (**b)** vs. cortical depth in all human patients.

**Supplementary Fig. 3.**
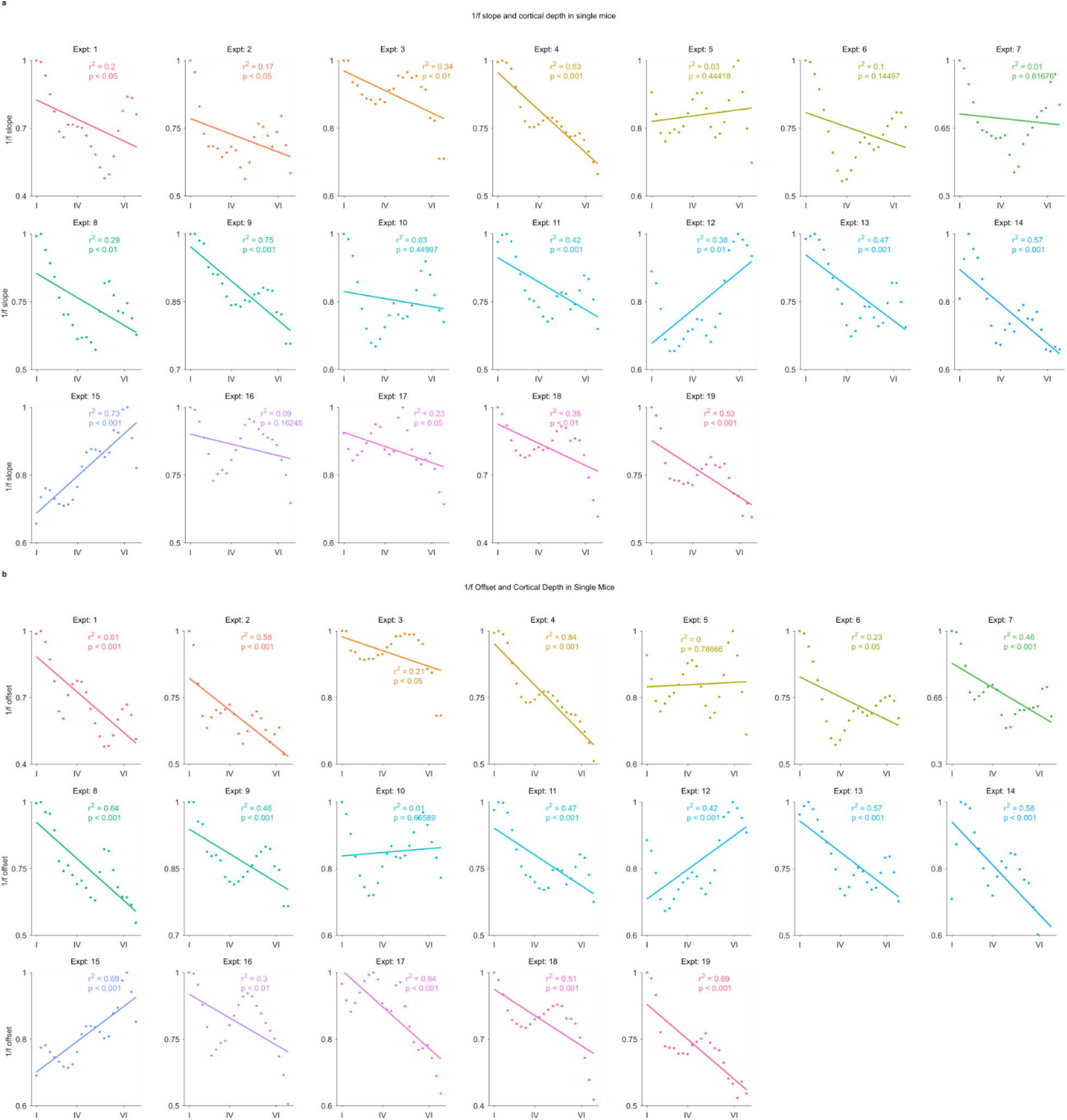
1/f slope and offset decrease with cortical depth in individual mouse experiments. 1/f slope (**a)** and offset (**b**) vs. cortical depth (x-axis) in all mouse experiments.

**Supplementary Fig. 4.**
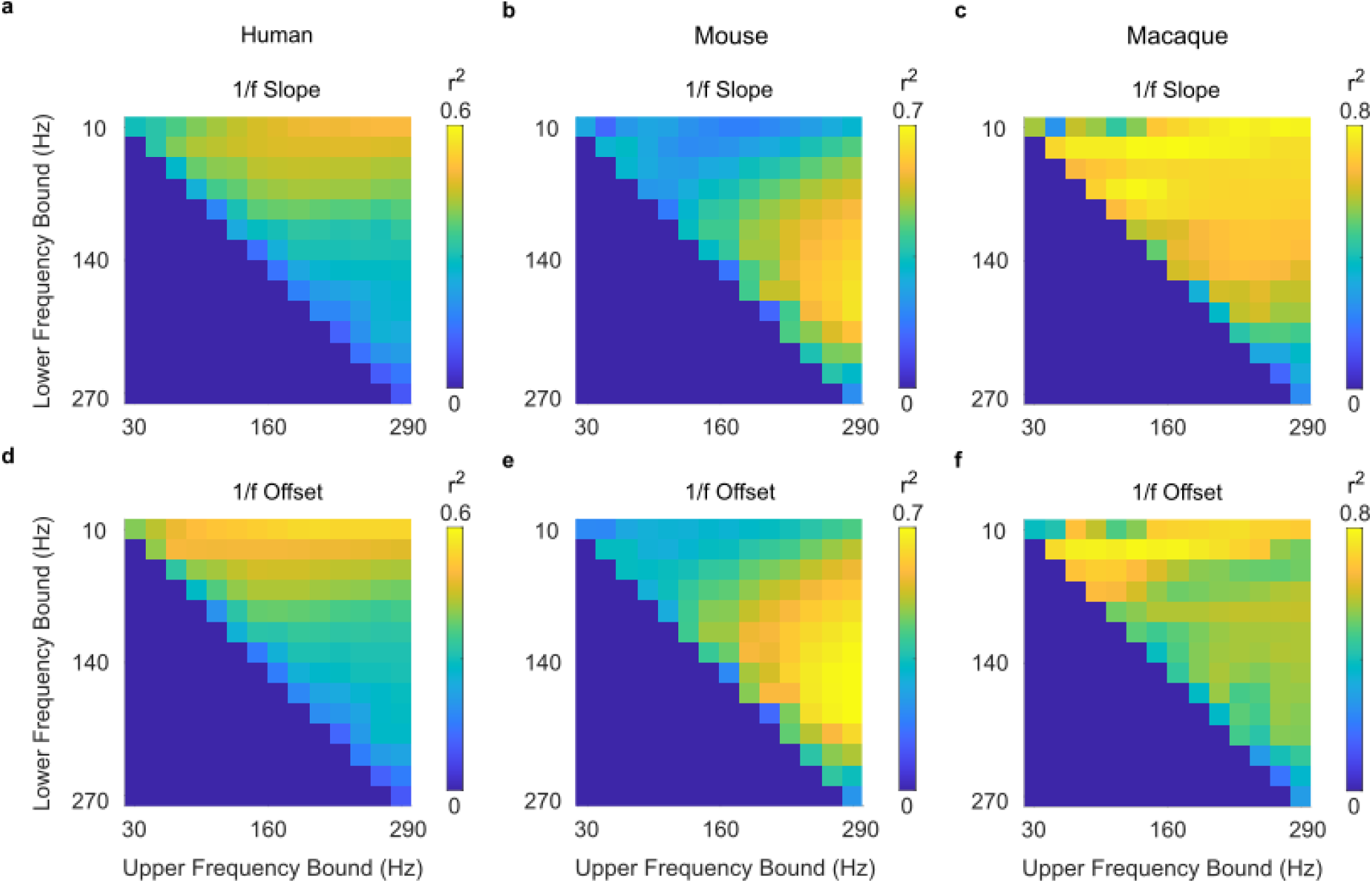
1/f slope and cortical depth are highly correlated across a wide range of frequency bands. For every frequency band between 10 and 270 Hz (20 Hz intervals), we found the correlation (r^2^) between 1/f slope or offset and depth within each patient (human) / experiment (mouse, macaque), and then averaged this value across patients / sessions.

**Supplementary Fig. 5.**
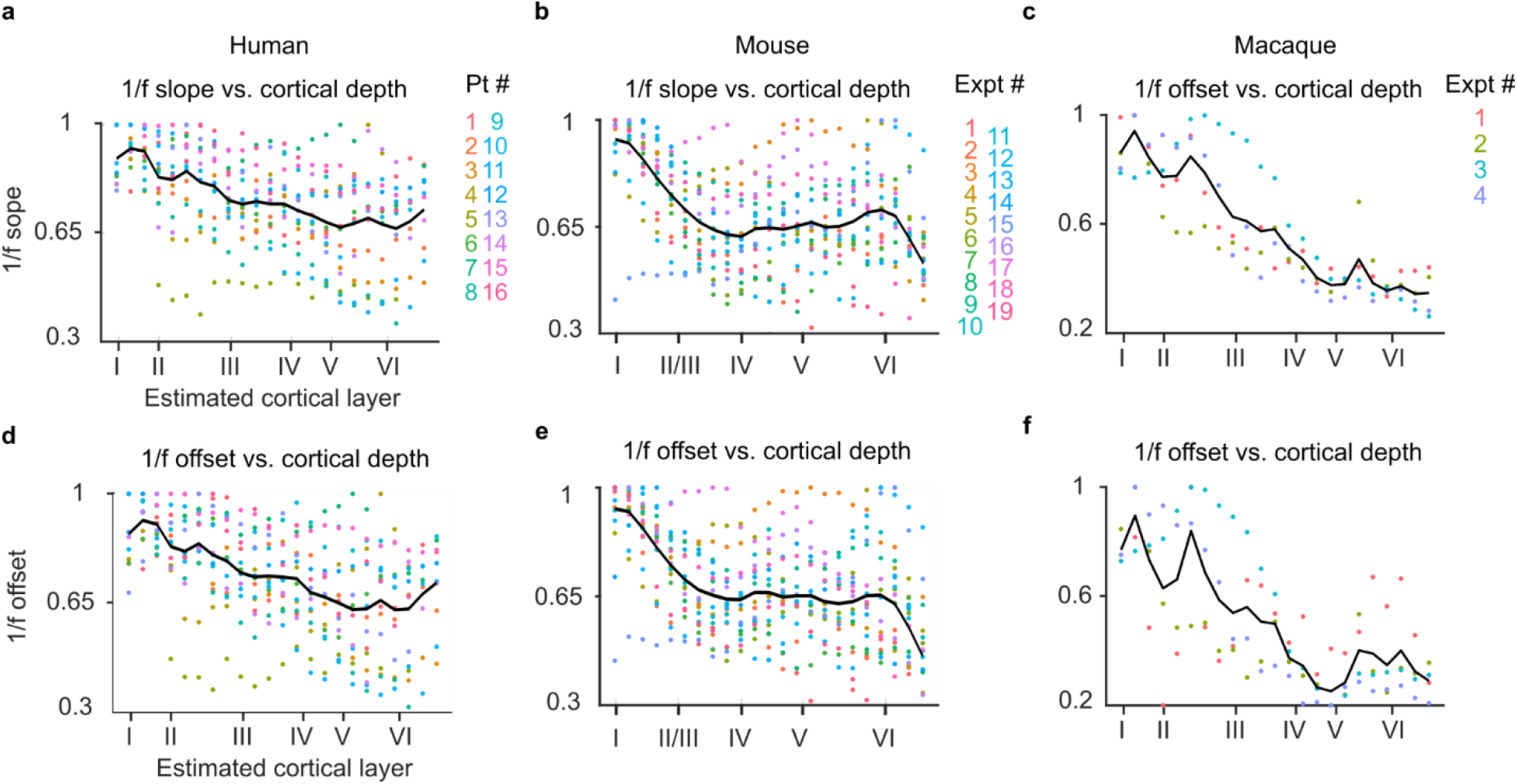
1/f slope flattens and offset decreases with cortical depth from 30-290 Hz. 1/f slope and offset vs. cortical depth in humans (**a, d**), mice (**b, e**) and macaques (**c, f**), fit using a frequency band of 30-290 Hz (as opposed to 1-290 Hz in our main results).

**Supplementary Fig. 6.**
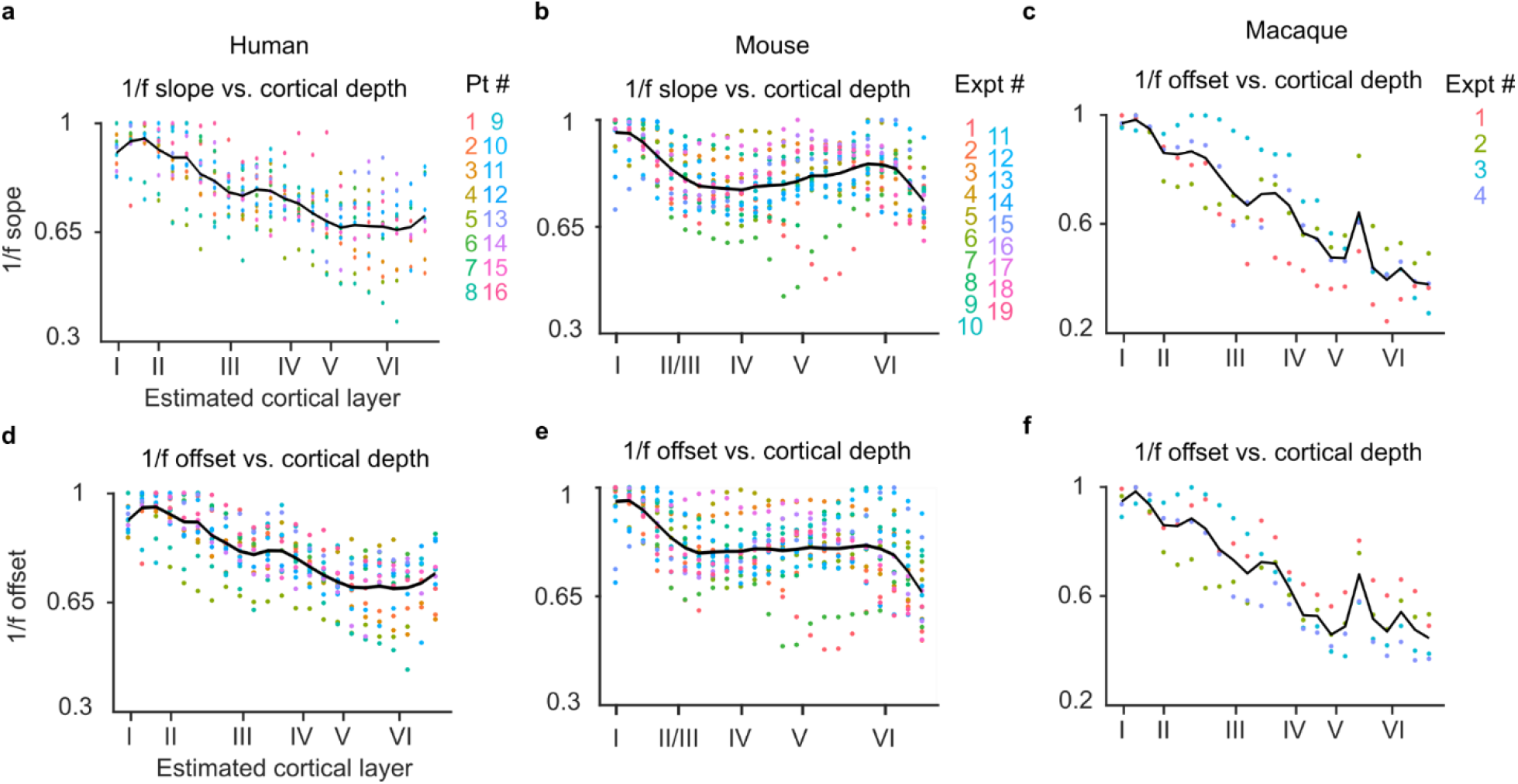
1/f slope and offsets fit with simple linear regression (polyfit.m) instead of the spectral parameterization algorithm yields similar results. 1/f slope and offsets across all humans (**a, d**), mice (**b, e**), and macaques (**c, f**) normalized within each experiment.

## References

Beaulieu-Laroche L, Toloza EHS, van der Goes MS, Lafourcade M, Barnagian D, Williams ZM, Eskandar EN, Frosch MP, Cash SS, Harnett MT. 2018. Enhanced Dendritic Compartmentalization in Human Cortical Neurons. Cell. doi:10.1016/j.cell.2018.08.045

Bédard C, Destexhe A. 2009. Macroscopic models of local field potentials and the apparent 1/f noise in brain activity. Biophys J. doi:10.1016/j.bpj.2008.12.3951

Bédard C, Kröger H, Destexhe A. 2006. Does the 1/f frequency scaling of brain signals reflect self-organized critical states? Phys Rev Lett. doi:10.1103/PhysRevLett.97.118102

Beggs JM, Plenz D. 2003. Neuronal Avalanches in Neocortical Circuits. J Neurosci. doi:10.1523/jneurosci.23-35-11167.2003

Blume WT. 2006. Drug effects on EEG. J Clin Neurophysiol 23:306–311. doi:10.1097/01.wnp.0000229137.94384.fa

Bódizs R, Szalárdy O, Horváth C, Ujma PP, Gombos F, Simor P, Pótári A, Zeising M, Steiger A, Dresler M. 2021. A set of composite, non-redundant EEG measures of NREM sleep based on the power law scaling of the Fourier spectrum. Sci Rep. doi:10.1038/s41598-021-81230-7

Buzsáki G, Anastassiou C a., Koch C. 2012. The origin of extracellular fields and currents — EEG, ECoG, LFP and spikes. Nat Rev Neurosci 13:407–420. doi:10.1038/nrn3241

Buzsáki G, Draguhn A. 2004. Neuronal Oscillations in Cortical Networks. Sci 304:1926–1929. doi:10.1126/science.1099745

Cash SS, Halgren E, Dehghani N, Rossetti AO, Thesen T, Wang C, Devinsky O, Kuzniecky R, Doyle W, Madsen JR, Bromfield E, Eross L, Halasz P, Karmos G, Csercsa R, Wittner L, Ulbert I. 2009. The Human K-Complex Represents an Isolated Cortical Down-State. Science (80-) 324:1084–1087. doi:10.1126/science.1169626

Chaudhuri R, He BJ, Wang XJ. 2018. Random recurrent networks near criticality capture the broadband power distribution of human ECoG dynamics. Cereb Cortex. doi:10.1093/cercor/bhx233

Colombo MA, Napolitani M, Boly M, Gosseries O, Casarotto S, Rosanova M, Brichant J-F, Boveroux P, Rex S, Laureys S, Massimini M, Chieregato A, Sarasso S. 2019. The spectral exponent of the resting EEG indexes the presence of consciousness during unresponsiveness induced by propofol, xenon, and ketamine. Neuroimage 189:631–644. doi:https://doi.org/10.1016/j.neuroimage.2019.01.024

Csercsa R, Dombovári B, Fabó D, Wittner L, Erőss L, Entz L, Sólyom A, Rásonyi G, Szűcs A, Kelemen A, Jakus R, Juhos V, Grand L, Magony A, Halász P, Freund TF, Maglóczky Z, Cash SS, Papp L, Karmos G, Halgren E, Ulbert I. 2010. Laminar analysis of slow wave activity in humans. Brain 133:2814–2829.

Dave S, Brothers TA, Swaab TY. 2018. 1/f neural noise and electrophysiological indices of contextual prediction in aging. Brain Res 1691:34–43. doi:https://doi.org/10.1016/j.brainres.2018.04.007

Donoghue T, Haller M, Peterson EJ, Varma P, Sebastian P, Gao R, Noto T, Lara AH, Wallis JD, Knight RT, Shestyuk A, Voytek B. 2020. Parameterizing neural power spectra into periodic and aperiodic components. Nat Neurosci. doi:10.1038/s41593-020-00744-x

Eickhoff SB, Rottschy C, Zilles K. 2007. Laminar distribution and co-distribution of neurotransmitter receptors in early human visual cortex. Brain Struct Funct 212:255–267. doi:10.1007/s00429-007-0156-y

Freeman WJ, Zhai J. 2009. Simulated power spectral density (PSD) of background electrocorticogram (ECoG). Cogn Neurodyn 3:97–103. doi:10.1007/s11571-008-9064-y

Gao R. 2016. Interpreting the electrophysiological power spectrum. J Neurophysiol. doi:10.1152/jn.00722.2015

Gao R, Peterson EJ, Voytek B. 2017. Inferring synaptic excitation/inhibition balance from field potentials. Neuroimage 158:70–78. doi:10.1016/j.neuroimage.2017.06.078

Gao R, Van den Brink RL, Pfeffer T, Voytek B. 2020. Neuronal timescales are functionally dynamic and shaped by cortical microarchitecture. Elife. doi:10.7554/eLife.61277

Haegens S, Barczak A, Musacchia G, Lipton ML, Mehta AD, Lakatos P, Schroeder CE. 2015. Laminar Profile and Physiology of the α Rhythm in Primary Visual, Auditory, and Somatosensory Regions of Neocortex. J Neurosci 35:14341–52. doi:10.1523/JNEUROSCI.0600-15.2015

Halgren E, Kaestner E, Marinkovic K, Cash SS, Wang C, Schomer DL, Madsen JR, Ulbert I. 2015. Laminar profile of spontaneous and evoked theta: Rhythmic modulation of cortical processing during word integration. Neuropsychologia 76:108–124. doi:10.1016/j.neuropsychologia.2015.03.021

Halgren M, Fabó D, Ulbert I, Madsen JR, Erőss L, Doyle WK, Devinsky O, Schomer D, Cash SS, Halgren E. 2018. Superficial Slow Rhythms Integrate Cortical Processing in Humans. Sci Rep 8:2055. doi:10.1038/s41598-018-20662-0

Halgren M, Ulbert I, Bastuji H, Fabó D, Erőss L, Rey M, Devinsky O, Doyle WK, Mak-McCully R, Halgren E, Wittner L, Chauvel P, Heit G, Eskandar E, Mandell A, Cash SS. 2019. The generation and propagation of the human alpha rhythm. Proc Natl Acad Sci 201913092. doi:10.1073/pnas.1913092116

Harnett MT, Magee JC, Williams SR. 2015. Distribution and function of HCN channels in the apical dendritic tuft of neocortical pyramidal neurons. J Neurosci. doi:10.1523/JNEUROSCI.2813-14.2015

He BJ, Zempel JM, Snyder AZ, Raichle ME. 2010. The temporal structures and functional significance of scale-free brain activity. Neuron 66:353–369. doi:10.1016/j.neuron.2010.04.020

He BYJ. 2014. Scale-free brain activity: past, present, and future. Trends Cogn Sci 18:480–487. doi:Doi 10.1016/J.Tics.2014.04.003

Hutsler JJ, Lee D-G, Porter KK. 2005. Comparative analysis of cortical layering and supragranular layer enlargement in rodent carnivore and primate species. Brain Res 1052:71–81. doi:10.1016/j.brainres.2005.06.015

Kajikawa Y, Schoeder E. 2012. How local is the local field potential? Neuron 72:847–858. doi:10.1016/j.neuron.2011.09.029.How

Kajikawa Y, Schroeder CE. 2015. Generation of field potentials and modulation of their dynamics through volume integration of cortical activity. J Neurophysiol 113:339–351. doi:10.1152/jn.00914.2013

Kalmbach B, Buchin A, Miller JA, Bakken TE, Hodge RD, Chong P, de Frates R, Dai K, Gwinn RP, Cobbs C, Ko AL, Ojemann JG, Silbergeld DL, Koch C, Anastassiou CA, Lein E, Ting JT. 2018. h-channels contribute to divergent electrophysiological properties of supragranular pyramidal neurons in human versus mouse cerebral cortex. bioRxiv.

Kandel ER, Schwartz J, Jessel TM, Siegelbaum SA, Hudspeth AJ. 1991. Principles of Neural Science, Fifth Edition, Elsevier.

Kiebel SJ, Daunizeau J, Friston KJ. 2008. A hierarchy of time-scales and the brain. PLoS Comput Biol. doi:10.1371/journal.pcbi.1000209

Kole MHP, Hallermann S, Stuart GJ. 2006. Single <em>I</em>hChannels in Pyramidal Neuron Dendrites: Properties, Distribution, and Impact on Action Potential Output. J Neurosci 26:1677LP–1687.

Lendner JD, Helfrich RF, Mander BA, Romundstad L, Lin JJ, Walker MP, Larsson PG, Knight RT. 2020. An electrophysiological marker of arousal level in humans. Elife. doi:10.7554/eLife.55092

Logothetis NK, Kayser C, Oeltermann A. 2007. In vivo measurement of cortical impedance spectrum in monkeys: implications for signal propagation. Neuron 55:809–23. doi:10.1016/j.neuron.2007.07.027

Manning JR, Jacobs J, Fried I, Kahana MJ. 2009. Broadband Shifts in Local Field Potential Power Spectra Are Correlated with Single-Neuron Spiking in Humans. J Neurosci 29:13613–13620. doi:10.1523/JNEUROSCI.2041-09.2009

Markov NT, Vezoli J, Chameau P, Falchier A, Quilodran R, Huissoud C, Lamy C, Misery P, Giroud P, Ullman S, Barone P, Dehay C, Knoblauch K, Kennedy H. 2014. Anatomy of hierarchy: Feedforward and feedback pathways in macaque visual cortex. J Comp Neurol 522:225–259. doi:10.1002/cne.23458

Miller KJ, Sorensen LB, Ojemann JG, Den Nijs M. 2009a. Power-law scaling in the brain surface electric potential. PLoS Comput Biol 5. doi:10.1371/journal.pcbi.1000609

Miller KJ, Zanos S, Fetz EE, Den Nijs M, Ojemann JG. 2009b. Decoupling the cortical power spectrum reveals real-time representation of individual finger movements in humans. J Neurosci. doi:10.1523/JNEUROSCI.5506-08.2009

Milotti E. 2002. 1/f Noise: A pedagogical review. Arxiv Phys.

Murray JD, Bernacchia A, Freedman DJ, Romo R, Wallis JD, Cai X, Padoa-Schioppa C, Pasternak T, Seo H, Lee D, Wang X-J. 2014. A hierarchy of intrinsic timescales across primate cortex. Nat Neurosci 17:1661–3. doi:10.1038/nn.3862

Ness T V, Remme MWH, Einevoll GT. 2018. h-type membrane current shapes the local field potential (LFP) from populations of pyramidal neurons. J Neurosci.

Oostenveld R, Fries P, Maris E, Schoffelen J-M. 2011. FieldTrip: Open Source Software for Advanced Analysis of MEG, EEG, and Invasive Electrophysiological Data. Comput Intell Neurosci 2011:1–9. doi:10.1155/2011/156869

Ouyang G, Hildebrandt A, Schmitz F, Herrmann CS. 2020. Decomposing alpha and 1/f brain activities reveals their differential associations with cognitive processing speed. Neuroimage 205:116304. doi:https://doi.org/10.1016/j.neuroimage.2019.116304

Podvalny E, Noy N, Harel M, Bickel S, Chechik G, Schroeder CE, Mehta AD, Tsodyks M, Malach R. 2015. A unifying principle underlying the extracellular field potential spectral responses in the human cortex. J Neurophysiol. doi:10.1152/jn.00943.2014

Robertson MM, Furlong S, Voytek B, Donoghue T, Boettiger CA, Sheridan MA. 2019. EEG power spectral slope differs by ADHD status and stimulant medication exposure in early childhood. J Neurophysiol. doi:10.1152/jn.00388.2019

Runyan CA, Piasini E, Panzeri S, Harvey CD. 2017. Distinct timescales of population coding across cortex. Nature. doi:10.1038/nature23020

Schaworonkow N, Voytek B. 2021. Longitudinal changes in aperiodic and periodic activity in electrophysiological recordings in the first seven months of life. Dev Cogn Neurosci. doi:10.1016/j.dcn.2020.100895

Shirhatti V, Borthakur A, Ray S. 2016. Effect of reference scheme on power and phase of the local field potential. Neural Comput. doi:10.1162/NECO_a_00827

Siegle JH, Jia X, Durand S, Gale S, Bennett C, Graddis N, Heller G, Ramirez TK, Choi H, Luviano JA, Groblewski PA, Ahmed R, Arkhipov A, Bernard A, Billeh YN, Brown D, Buice MA, Cain N, Caldejon S, Casal L, Cho A, Chvilicek M, Cox TC, Dai K, Denman DJ, de Vries SEJ, Dietzman R, Esposito L, Farrell C, Feng D, Galbraith J, Garrett M, Gelfand EC, Hancock N, Harris JA, Howard R, Hu B, Hytnen R, Iyer R, Jessett E, Johnson K, Kato I, Kiggins J, Lambert S, Lecoq J, Ledochowitsch P, Lee JH, Leon A, Li Y, Liang E, Long F, Mace K, Melchior J, Millman D, Mollenkopf T, Nayan C, Ng L, Ngo K, Nguyen T, Nicovich PR, North K, Ocker GK, Ollerenshaw D, Oliver M, Pachitariu M, Perkins J, Reding M, Reid D, Robertson M, Ronellenfitch K, Seid S, Slaughterbeck C, Stoecklin M, Sullivan D, Sutton B, Swapp J, Thompson C, Turner K, Wakeman W, Whitesell JD, Williams D, Williford A, Young R, Zeng H, Naylor S, Phillips JW, Reid RC, Mihalas S, Olsen SR, Koch C. 2021. Survey of spiking in the mouse visual system reveals functional hierarchy. Nature. doi:10.1038/s41586-020-03171-x

Spitmaan M, Seo H, Lee D, Soltani A. 2020. Multiple timescales of neural dynamics and integration of task-relevant signals across cortex. Proc Natl Acad Sci U S A. doi:10.1073/pnas.2005993117

Steinmetz NA, Zatka-Haas P, Carandini M, Harris KD. 2019. Distributed coding of choice, action and engagement across the mouse brain. Nature. doi:10.1038/s41586-019-1787-x

Stock A-K, Pertermann M, Mückschel M, Beste C. 2019. High-dose ethanol intoxication decreases 1/f neural noise or scale-free neural activity in the resting state. Addict Biol n/a:e12818. doi:10.1111/adb.12818

Timmermann C, Roseman L, Schartner M, Milliere R, Williams LTJ, Erritzoe D, Muthukumaraswamy S, Ashton M, Bendrioua A, Kaur O, Turton S, Nour MM, Day CM, Leech R, Nutt DJ, Carhart-Harris RL. 2019. Neural correlates of the DMT experience assessed with multivariate EEG. Sci Rep 9:16324. doi:10.1038/s41598-019-51974-4

Trongnetrpunya A, Nandi B, Kang D, Kocsis B, Schroeder CE, Ding M. 2016. Assessing Granger Causality in Electrophysiological Data: Removing the Adverse Effects of Common Signals via Bipolar Derivations. Front Syst Neurosci 9. doi:10.3389/fnsys.2015.00189

Ulbert István, Halgren E, Heit G, Karmos G. 2001. Multiple microelectrode-recording system for human intracortical applications. J Neurosci Methods 106:69–79. doi:10.1016/S0165-0270(01)00330-2

Ulbert Istvan, Karmos G, Heit G, Halgren E. 2001. Early discrimination of coherent versus incoherent motion by multiunit and synaptic activity in human putative MT+. Hum Brain Mapp 13:226–238. doi:10.1002/hbm.1035

V. Stewart C, Plenz D. 2006. Inverted-U Profile of Dopamine-NMDA-Mediated Spontaneous Avalanche Recurrence in Superficial Layers of Rat Prefrontal Cortex. J Neurosci 26:8148–8159. doi:10.1523/JNEUROSCI.0723-06.2006

Veerakumar A, Tiruvadi V, Howell B, Waters AC, Crowell AL, Voytek B, Riva-Posse P, Denison L, Rajendra JK, Edwards JA, Bijanki KR, Choi KS, Mayberg HS. 2019. Field potential 1/f activity in the subcallosal cingulate region as a candidate signal for monitoring deep brain stimulation for treatment-resistant depression. J Neurophysiol. doi:10.1152/jn.00875.2018

Voytek B, Kramer MA, Case J, Lepage KQ, Tempesta ZR, Knight RT, Gazzaley A. 2015. Age-Related Changes in 1/f Neural Electrophysiological Noise. J Neurosci 35:13257–13265. doi:10.1523/JNEUROSCI.2332-14.2015

Wang Q, Ding SL, Li Y, Royall J, Feng D, Lesnar P, Graddis N, Naeemi M, Facer B, Ho A, Dolbeare T, Blanchard B, Dee N, Wakeman W, Hirokawa KE, Szafer A, Sunkin SM, Oh SW, Bernard A, Phillips JW, Hawrylycz M, Koch C, Zeng H, Harris JA, Ng L. 2020. The Allen Mouse Brain Common Coordinate Framework: A 3D Reference Atlas. Cell. doi:10.1016/j.cell.2020.04.007

Waschke L, Donoghue T, Fiedler L, Smith S, Garrett DD, Voytek B, Obleser J. 2021. Modality-specific tracking of attention and sensory statistics in the human electrophysiological spectral exponent. bioRxiv.

Zilles K, Palomero-Gallagher N. 2017. Multiple transmitter receptors in regions and layers of the human cerebral cortex. Front Neuroanat. doi:10.3389/fnana.2017.00078

